# Cladistic hypotheses as degree of equivalence relational structures: implications for three-item statements

**DOI:** 10.1101/2021.01.14.426769

**Authors:** Valentin Rineau, Stéphane Prin

**Author notes:** These authors contributed equally to this work.

## Abstract

Three-item statements, as minimal informative rooted binary phylogenetic trees on three items, are the minimal units of cladistic information. Their importance for phylogenetic reconstruction, consensus and supertree methods relies on both (i) the fact that any cladistic tree can always be decomposed into a set of three-item statements, and (ii) the possibility, at least under some conditions, to build a new cladistic tree by combining all or part of the three-item statements deduced from several prior cladistic trees. In order to formalise such procedures, several *k*-adic rules of inference, i.e., rules that allow us to deduce at least one new three-item statement from exactly *k* other ones, have been identified. However, no axiomatic background has been proposed, and it remains unknown if a particular *k*-adic rule of inference can be reduced to more basic rules. In order to solve this problem, we propose here to define three-item statements in terms of degree of equivalence relations. Given both the axiomatic definition of the latter and their strong connection to hierarchical classifications, we establish a list of the most basic properties for three-item statements. With such an approach, we show that it is possible to combine five three-item statements from basic rules although they are not combinable only from dyadic rules. Such a result suggests that all higher *k*-adic rules are well reducible to a finite set of simpler rules.

## 1 Introduction

The increase in biological data as well as in processing capacities has made it possible to generate more and more phylogenetic trees on various taxonomic groups. New problems have arisen with the increasing number of trees concerning the manipulation and amalgamation of several trees into a unified hypothesis in order to make their multitude intelligible, and, with them, new phylogenetic methods of tree reconstruction and combination (Bininda-Edmonds 2004). Supertree-like methods, including species tree estimation under the multispecies coalescent (e.g., ASTRAL; Zhang et al. 2018) have become central for phylogeny estimation in the past ten years. As a result, a growing number of algorithms have been designed to solve supertree-like problems (Nguyen et al. 2012; Sevillya et al. 2016; Fleischauer and Böcker 2017). The development of these methods and practices implies more than ever to have a relevant formal description of phylogenetic trees as mathematical structures on which biologists can rely.

Among the first to investigate the combinatorial properties of hierarchical *n*-trees and/or unrooted *n*-trees, Dobson (1974) introduced a symbolism for quartets (i.e., minimal informative unrooted subtrees). Using the four-points conditions, she then deduced as corollaries two inference rules for quartets. After that, Dekker (1986) proposed a more systematic approach to inference rules for quartets. She introduced the concept of *k*-adic rules of inference, i.e., rules such that given *k* quartets, at least one other follows. She then proved the existence of not only two dyadic rules of inference, but also several higher-order *k*-adic rules of inference irreducible to the former. Dekker eventually conjectured from this result that such higher-order rules actually are infinite. Subsequently, Bryant and Steel (1995) transposed Dekker’s work to three-item statements (minimal informative rooted subtrees; 3is hereafter) and proved Dekker’s conjecture for both 3is and quartets. Note that their approach, although not axiomatic, is formal and abstract. Using a non-formal and example-based approach, Wilkinson et al. (2004) finally inferred three dyadic rules for 3is from the two dyadic rules for quartets proposed by Dekker and explicitly interpreted these rules as dependency rules. These works show that understanding the problem of amalgaming larger trees from combination of subtrees as a problem of *k*-adic rules lead to an infinite complexity, that is to say, there are an infinite quantity of higher-order rules of combination irreducible to a finite set of dyadic rules of combination.

Interestingly, none of the previous work is axiomatic, i.e., none of them consists to start from a consistent, necessary, and sufficient primitive rules from which all the other rules could be deduced. As a result, it is hard to determine whether or not the higher rules really are irreducible to more basic rules. Especially, one cannot determine whether the fact that higher order rules are irreducible to dyadic rules means that these higher-order rules are irreducible to more basic rules. In other words, it is hard to determine whether dyadic rules are the only possible most basic rules. In this paper, we propose to address this issue for the specific case of rooted trees and 3is with a different framework. On the one hand we propose to define 3is in terms of specific ternary relations on a set called “degree of equivalence relations”. The idea here is to rely on both the axiomatic properties of these relations and their bijective connexion to hierarchical classifications. On the other hand, we propose to illustrate the fecundity of such an approach by considering an example of five compatible but pairwise non-combinable 3is (Wilkinson pers. com.).

Usually, 3is are encountered in cladistic phylogenetics. Indeed, the latter consists of establishing the degree of kinship relationship *sensu* Prin (2016), also called “phylogenetic relationship” (Hennig 1965; Nelson 1994), “relationship of common ancestry” (Nelson 1970, 1972, 1973), or “cladistic relationship” (Camin and Sokal 1965; Hull 1970) between the *n* taxonomic units^1^ (TU hereafter) of a given set. The degree of kinship relationship is a ternary biological relationship of kinship that consists to say that given three biological objects of the same kind (organisms or taxa), two of them are closely related to each other than either is to the third. But any 3is is precisely an assertion that, given three distinct objects of a same kind (e.g., TU), states that two of them are closely related to each other than either is to the third. Hence the possibility to deduce a consistent set of 3is from any cladistic tree, i.e., any hierarchical *n*-tree formalising a given cladistic hypothesis. By accepting that degree of equivalence relations formalise the degree of kinship relationship, that suggests the possibility to formally define 3is in terms of degree of equivalence relations.

In order to achieve this aim, this paper is structured in the following seven steps: (i) we provide an axiomatic definition of degree of equivalence relations, (ii) we recall the axiomatic definition of hierarchical classifications, (iii) we address the mathematical equivalence between degree of equivalence relations and hierarchical classifications, (iv) we draw the consequences on the mathematical structures of cladistic hypotheses, (v) we define 3is in terms of degree of equivalence relations and investigate some consequences of it, (vi) we generalise some basic rules seen before to higher-order rules, and then (vii) we apply these consequences for a specific set of compatible but pairwise non-combinable 3is.

## 2 Degree of equivalence relation

Let ⟨*A, T* ⟩ be an ordered pair. In this case, ⟨*A, T* ⟩ is also a *degree of equivalence relational structure* if and only if (a) *A* is a non-empty set and (b) *T* is a *degree of equivalence relation* on *A*, i.e., a ternary relation on *A* (*T* ⊆ *A*^3^) such that for all *w, x, y, z* ∈ *A*:^2^

1. (*x, y, z*) ∈ *T* → [(*y, x, z*) ∈ *T* ∧ (*x, z, y*) ∉*T*] (1-2 symmetry & 2-3 asymmetry)
2. *x* ≠*y* → (*x, x, y*) ∈ *T* (1-2 conditional reflexivity)
3. [(*x, y, w*) ∈ *T* ∧ (*y, z, w*) ∈ *T*] → (*x, z, w*) ∈ *T* (1-2 transitivity)
4. (*w, x, z*) ∈ *T* → [(*w, x, y*) ∈ *T* ∨ (*w, y, z*) ∈ *T*] (2-3 negative transitivity) Interestingly, these axiomatic properties imply several derived properties. Below are some of them (here too, for all *w, x, y, z* ∈ *A*):
5. ⟨*x, y, z* ⟩ ∈ *T* → ⟨*z, y, x* ⟩ ∉*T* (1-3 asymmetry)
6. ⟨*x, y, x* ⟩ ∉*T* ∧ ⟨*x, y, y*⟩ ∉*T* (1-3 & 2-3 anti-reflexivity)
7. ⟨*x, x, y*⟨ ∈ *T* ∧ *x* ≠ *y* (1-2 relative identity)
8. ⟨*x, x, x* ⟩ ∉*T* (anti-reflexivity)
9. [⟨*w, x, y*⟩ ∈ *T* ∧ ⟨*w, y, z* ⟩ ∈ *T*] → [⟨*w, x, z* ⟩ ∈ *T* ∧ ⟨*x, y, z* ⟩ ∈ *T*] (2-3 enhanced transitivity)
10. [⟨*x, w, y*⟩ ∈ *T* ∧ ⟨*y, w, z* ⟩ ∈ *T*] → [⟨*x, w, z* ⟩ ∈ *T* ∧ ⟨*x, y, z* ⟩ ∈ *T*] (1-3 enhanced transitivity)
11. [⟨*w, x, y*⟩ ∈ *T* ∧ ⟨*x, y, z* ⟩ ∈ *T*] → [⟨*w, x, z* ⟩ ∈ *T* ∧ ⟨*w, y, z* ⟩ ∈ *T*] (crossed transitivity)
12. [⟨*w, x, y*⟩ ∈ *T* ∧ ⟨*y, z, w* ⟩ ∈ *T*] → [⟨*w, x, z* ⟩ ∈ *T* ∧ ⟨*y, z, x* ⟩ ∈ *T*] (pair separation)

Hereafter, the set *A* will be considered only as finite. Now, let *𝒟*_*A*_ be the set of all degree of equivalence relations on *A*. Then let *T* ∈ *𝒟*_*A*_ and let ⟨*x, y, z* ⟩ ∈ *T*. In this case, ⟨*x, y, z* ⟩ is trivial (or non-informative) if *x* = *y* and non-trivial (or informative) if *x* ≠ *y*. It follows a single trivial degree of equivalence relation *T*_*A*0_ on *A* such that

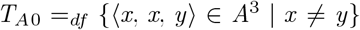

i.e.,

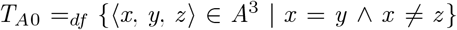

Given the properties above, it is not hard to see that *T* ⊇ *T*_*A*0_ for any *T* ∈ *𝒟*_*A*_. In corollary, any *T* ∈ *𝒟*_*A*_ is non-trivial if and only if *T* ⊃ *T*_*A*0_. Let us now define the set

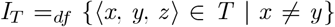

of all non-trivial 3-tuples of *T*. In this case, one has (i) *I*_*T*_ *∉𝒟*_*A*_, (ii) *T* = *T*_*A*0_ ∪ *I*_*T*_, and (iii) *T*_*A*0_ ∩ *I*_*T*_ = ∅.

To conclude, note that the symmetry-based properties of *T* imply that there are always three incompatible pairs of non-trivial 3-tuples for all distinct *x, y, z* ∈ *A*: {⟨*x, y, z* ⟩, ⟨*y, x, z* ⟩}, {⟨*x, z, y*⟩, ⟨*z, x, y*⟩}, and {⟨*y, z, x* ⟩, ⟨*z, y, x* ⟩}.

In other words, either {⟨*x, y, z* ⟩, ⟨*y, x, z* ⟩} ⊆ *T* and {⟨*x, z, y*⟩, ⟨*z, x, y*⟩, ⟨*y, z, x* ⟩, ⟨*z, y, x* ⟩} ∩ *T* = ∅, or {⟨*x, z, y*⟩, ⟨*z, x, y*⟩} ⊆ *T* and {⟨*x, y, z* ⟩, ⟨*y, x, z* ⟩, ⟨*y, z, x* ⟩, ⟨*z, y, x* ⟩} ∩ *T* = ∅, or {⟨*y, z, x* ⟩, ⟨*z, y, x* ⟩} ⊆ *T* and {⟨*x, y, z* ⟩, ⟨*y, x, z* ⟩, ⟨*x, z, y*⟩, ⟨*z, x, y*⟩} ∩ *T* = ∅, or {⟨*x, y, z* ⟩, ⟨*y, x, z* ⟩, ⟨*x, z, y*⟩, ⟨*z, x, y*⟩, ⟨*y, z, x* ⟩, ⟨*z, y, x* ⟩} ∩ *T* = ∅.

## 3 Hierarchical classification

Let ⟨*A, H* ⟩ be an ordered pair. In this case, ⟨*A, H* ⟩ is a finite *hierarchical classificatory structure* if and only if (a) *A* is a finite non-empty set, and (b) *H* is a *hierarchical classification*^3^ of *A*, i.e., a subset of *℘*(*A*) \ {∅} such that

13. *H* ⊇ {*X* ⊆ *A* | *X* = *A* ∨ |*X* | = 1}

14. (∀*X, Y* ∈ *H*) (*X* ∩ *Y* = ∅ ∨ *X* ⊆ *Y* ∨ *Y* ⊆ *X*)

Hereafter, the elements of *H* will be called its “*clusters* “. Now, let *ℋ* _*A*_ be the set of all possible hierarchical classifications of *A*. Then let *H* ∈ *ℋ* _*A*_ and *X* ∈ *H*. In this case, *X* is trivial (or non-informative) if *X* = *A* and/or |*X* | = 1, and non-trivial (or informative) if 2 ≤ |*X* | *<* |*A*|. Any non-empty set *A* thus implies one and only one trivial hierarchical classification *H*_*A*0_ of *A* such that

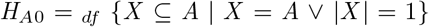

i.e.,

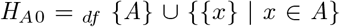

Here too, the properties above imply *H* ⊇ *H*_*A*0_ for any *H* ∈ *H* _*A*_. In corollary, any *ℋ* ∈ *H* _*A*_ is non-trivial if and only if *H* ⊃ *ℋ*_*A*0_. Then considers the set

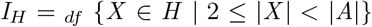

of all non-trivial clusters of *H*. In this case, (i) *I*_*H*_ *∉ℋ* _*A*_, (ii) *H* = *H*_*A*0_ ∪ *I*_*H*_, and (iii) *H*_*A*0_ ∩ *I*_*H*_ = ∅.

To conclude, note that property 14 implies an incompatibility between the possible non-trivial clusters of *H* analogous to the incompatibility between the non-trivial 3-tuples of *T*. Indeed, suppose for all distinct *x, y, z* ∈ *A* and any *X* ∈ *H* that there is at least one *Y* ∈ *H* such that {*x, y*} ⊆ *Y* and *z ∉Y*. In this case, *X* ⊃ *Y* if either {*x, z*} ⊆ *X* or {*y, z*} ⊆ *X*.

## 4 Degree of equivalence relation and hierarchical classification

Remarkably, there is a mathematical equivalence between (the) degree of equivalence relations (on *A*) and (the) hierarchical classifications (of *A*) (Colonius and Schulze 1981; McMorris and Powers 2003). This equivalence can be described as follows (Prin 2016: 2^*nd*^ appendix). (1) Let *T* ∈ *𝒟*_*A*_ and {*x, y*} ⊆ *A*. Then, define the *T*-relative *degree of equivalence class* of *x* and *y* as the set [*x* & *y*]_*T*_ of the elements *z* of *A* such that *x* and *y* are not *T*-equivalent compared to *z*, i.e.,

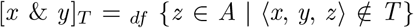

In this case, the *T*-relative *propinquity set* of *A*, i.e., the set

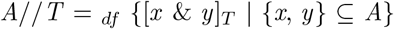

i.e.,

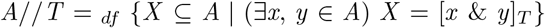

of all possible degree of equivalence classes induced by *T* is necessarily a hierarchical classification of *A*. Hence, there is a function *f*: *𝒟*_*A*_ → *ℋ* _*A*_ such that *f* (*T*) = *A*//*T* for any *T* ∈ *D*_*A*_. (2) Let *H* ∈ *ℋ* _*A*_. In this case, the ternary relation *T*_*H*_ on *A* such that for any {*x, y, z*}, ⟨*x, y, z* ⟩ ∈ *T*_*H*_ if and only if (i) ⟨*x, y, z* ⟩ ∈ *A*^3^ and (ii) there is at least one *X* ∈ *H* containing *x* and *y* but not *z*, i.e.,

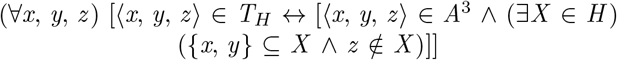

i.e.,

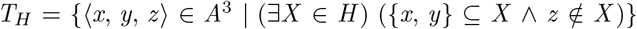

is necessarily a degree of equivalence relation on *A*. Thus, there is a function *g*: *ℋ* _*A*_ → *𝒟*_*A*_ such that *g*(*H*) = *T*_*H*_ for any *H* ∈ *ℋ* _*A*_. (3) Let (*T, H*) ∈ *𝒟*_*A*_ × *ℋ* _*A*_. In this case, not only *g*(*f* (*T*)) = *T*, but also *f* (*g*(*H*)) = *H*. Hence, *f* and *g* are both injective (which implies both |*ℋ*_*A*_| ≤ |*𝒟* _*A*_| and |*𝒟* _*A*_| ≤ |*ℋ*_*A*_|) and so, bijective too (since |*ℋ*_*A*_| = |*𝒟* _*A*_|). (4) Finally, let ⟨*T, H* ⟩ ∈ *D*_*A*_ × *H* _*A*_ again. The preceding implying that *f* (*T*) = *H* if and only if g(*H*) = *T*, it follows that *f* and *g* are converses to each other. Degree of equivalence relations and hierarchical classifications are therefore just two ways for saying that given three objects, either two of them are equivalent compared to the third, or there is no possibility to separate one of them from the other two. In particular, *T*_*A*0_ and *H*_*A*0_ correspond to each other, i.e., *f* (*T*_*A*0_) = *H*_*A*0_ and *g*(*H*_*A*0_) = *T*_*A*0_.

## 5 Consequences on cladistic hypotheses

Following Prin (2016), we adopt the view according to which any phylogenetic hypothesis can basically be formalised as a finite simple *k*-ary relational structure, i.e., an ordered pair

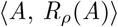

such that: (a) *A* is a finite non-empty set of mutually disjoint TU; and (b) *R*_*ρ*_(*A*) is a *k*-ary relation on *A*, i.e., a subset of *A*^*k*^ such as

a. *R*_*ρ*_(*A*) = {⟨*x* _1_, *x* _2_, …, *x*_*k*_⟩ ∈ *A*^*k*^ | *ρ*(*x* _1_, *x* _2_, …, *x*_*k*_)}, and
b. the *k*-adic propositional function “*ρ*(*x* _1_, *x* _2_, …, *x*_*k*_)” corresponds to the complete predicative formulation of a given *k*-ary concept of kinship in our language (e.g., “*x* is an ancestor of *y*” for the ancestor-descendant relationship).^4^

Such a general schema can easily be applied to cladistic phylogenetic hypotheses, i.e., phylogenetic hypotheses expressing the degree of kinship relationship between the TU of a given set. For this, one needs both (i) a predicative formulation of the degree of kinship relationship, and (ii) to know its formal properties. For the predicative formulation, let us recall that this relationship of kinship is a ternary one that consists to say that given three biological entities (organisms and their parts or taxa and their properties), two of them are more closely related than either is to the third (or, if you prefer, that two of them are closely related in respect to the third). Thus, possible propositional functions can be either “*x* is more closely related to *y* than to *z* “or “*x* is (closely) related to *y* compared to *z* “, abbreviated “*δ*(*x, y, z*)” hereafter. As for its formal properties, it has been argued in Prin (2016) that this relationship mathematically corresponds to degree of equivalence relations. Consequently, any cladistic phylogenetic hypothesis *D* can basically be formalised as a finite non-trivial degree of equivalence relational structure, i.e., an ordered pair

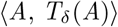

such that: (a) *A* is a finite non-empty set of at least three disjoint TU (i.e., |*A*| ≥ 3); and (b) *T*_*δ*_(*A*) is a non-trivial degree of equivalence relation on *A*, a subset of *A*^3^ such as:

a. *T*_*δ*_(*A*) = {⟨*x, y, z* ⟩ ∈ *A*^3^ | *δ*(*x, y, z*)}, and
b. there are at least three distinct *x, y, z* ∈ *A* such that either {⟨*x, y, z* ⟩, ⟨*y, x, z* ⟩} ⊆ *T* and {⟨*x, z, y*⟩, ⟨*z, x, y*⟩, ⟨*y, z, x* ⟩, ⟨*z, y, x* ⟩} ∩ *T* = ∅, or {⟨*x, z, y*⟩, ⟨*z, x, y*⟩} ⊆ *T* and {⟨*x, y, z* ⟩, ⟨*y, x, z* ⟩, ⟨*y, z, x* ⟩, ⟨*z, y, x* ⟩} ∩ *T* = ∅, or {(*y, z, x*), (*z, y, x*)} ⊆ *T* and {⟨*x, y, z* ⟩, ⟨*y, x, z* ⟩, ⟨*x, z, y*⟩, ⟨*z, x, y*⟩} ∩ *T* = ∅.

From this first formalisation follow at least three other ones. Indeed, if *D* is formalizable as ⟨*A, T*_*δ*_(*A*) ⟩, it is then formalizable as a finite hierarchical classificatory structure ⟨*A, H*_*δ*_(*A*) ⟩ such that *H*_*δ*_(*A*) = *A*//*T*_*δ*_(*A*). But if true, that also means that we can formalise *D* as a finite (binary) relational structure of strict order ⟨*H*_*δ*_(*A*), *R*_⊃_[*H*_*δ*_(*A*)] ⟩ such that

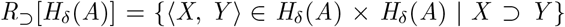

And finally, the latter implies the possibility to formalise *D* as a covering relational structure, i.e., an ordered pair ⟨*H*_*δ*_(*A*), *R*_⊐_[*H*_*δ*_(*A*)] ⟩ such that the covering relation *R*_⊐_[*H*_*δ*_(*A*)] is nothing but the transitive reduction of *R*_⊃_[*H*_*δ*_(*A*)], i.e., the greatest anti-transitive binary relation on *H*_*δ*_(*A*) included in *R*_⊃_[*H*_*δ*_(*A*)]. Note that the latter can be reinterpreted as a simple directed graph ⟨*V, E* ⟩ where each vertex *v* ∈ *V* bijectively corresponds to one cluster *X* ∈ *H*_*δ*_(*A*), and each directed edge *e* ∈ *E* bijectively corresponds to one ordered pair ⟨*X, Y* ⟩ ∈ *R*_⊐_[*H*_*δ*_(*A*)] (i.e., one immediate proper inclusion link *X* ⊐ *Y* between the two distinct clusters *X, Y* ∈ *H*_*δ*_(*A*)). But *R*_⊃_[*H*_*δ*_(*A*)] is also a fully ramified hierarchical strict order relation as it satisfies three important properties (other than anti-reflexivity and transitivity): (i) it has a minimum element (i.e., a single origin), which is its exhaustive cluster (*A* itself); (ii) it has no convergent ramification (i.e., no reticulation); and (iii) it has a complete divergent ramification. As a result, ⟨*H*_*δ*_(*A*), *R*_⊐_[*H*_*δ*_(*A*)] ⟩ is also a hierarchical *n*-tree, i.e., a directed tree such that its single root, its *n* leaves, and its (internal) nodes respectively (and bijectively) corresponds to the exhaustive cluster of *H*_*δ*_(*A*) (*A* itself), the *n* terminal clusters of *H*_*δ*_(*A*) (the *n* singletons {*x*} such that *x* ∈ *A*), and the non-trivial clusters of *H*_*δ*_(*A*). As we shall see below, this last formalisation justifies why one can represent cladistic hypotheses by dendrograms.

We can now illustrate this approach with a very simple theoretical example. Let *A* = {*a, b, c, d*} be a set of four distinct TU and let

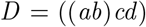

be a *A*-relative cladistic hypothesis saying that *a* and *b* are more closely related than either is to *c* and *d* (or, if you prefer, saying that *a* and *b* are closely related compared to *c* and *d*). In this case, the application of “*δ*(*x, y, z*)” into *A* (i.e., the searching for the 3-tuples (*x, y, z*) ∈ *A*^3^ for which “*δ*(*x, y, z*)” is true) implies the following statements (and nothing else): “*δ*(*a, a, b*)”, “*δ*(*a, a, c*)”, “*δ*(*a, a, d*)”, “*δ*(*b, b, a*)”, “*δ*(*b, b, c*)”, “*δ*(*b, b, d*)”, “*δ*(*c, c, a*)”, “*δ*(*c, c, b*)”, “*δ*(*c, c, d*)”, “*δ*(*d, d, a*)”, “*δ*(*d, d, b*)”, “*δ*(*d, d, c*)”, “*δ*(*a, b, c*)”, “*δ*(*b, a*, “, “*δ*(*a, b, d*)”, “*δ*(*b, a, d*)”. But given that *T*_*δ*_(*A*) = {(*x, y, z*) ∈ *A*^3^ | *δ*(*x, y, z*)}, it follows that

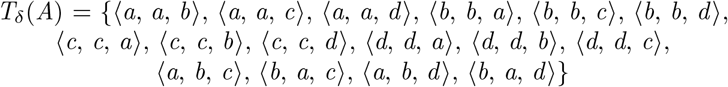

i.e.,

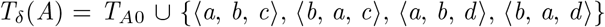

Consequently, *T*_*δ*_(*A*) is a non-trivial degree of equivalence relation on *A* and so, ⟨*A, T*_*δ*_(*A*) ⟩ is well a finite non-trivial degree of equivalence relational structure. Hence our first (and more basic) formalisation of *D*.

Then, let us recall that for all *x, y* ∈ *A*, the *T*_*δ*_(*A*)-relative degree of equivalence class of *x* and *y* is the set [*x* & *y*]_*T*_δ (*A*) = _*df*_ {*z* ∈ *A* | (*x, y, z*) ∈*/ T*_*δ*_(*A*)}. One therefore obtains: [*a* & *a*]_*T*_δ (*A*) = {*a*}, [*b* & *b*]_*T*_δ (*A*) = {*b*}, [*c* & *c*]_*T*_δ (*A*) = {*c*}, [*d* & *d*]_*T*_δ (*A*) = {*d*}, [*a* & *c*]_*T*_δ (*A*) = [*c* & *a*]_*T*_δ (*A*) = [*a* & *d*]_*T*_δ (*A*) = [*d* & *a*]_*T*_δ (*A*) = [*b* & *c*]_*T*_δ (*A*) = [*c* & *b*]_*T*_δ (*A*) = [*b* & *d*]_*T*_δ (*A*) = [*d* & *b*]_*T*_δ (*A*) = [*c* & *d*]_*T*_δ (*A*) = [*d* & *c*]_*T*_δ (*A*) = {*a, b, c, d*}, and [*a* & *b*]_*T*_δ (*A*) = [*b* & *a*]_*T*_δ (*A*) = {*a, b*}. But given that *A*//*T*_*δ*_(*A*) = _*df*_ {*X* ⊆ *A* | (∃*x, y* ∈ *A*) *X* = [*x* & *y*]_*T*_δ (*A*)}, it follows that

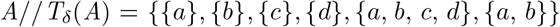

i.e.,

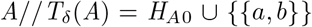

By defining *H*_*δ*_(*A*) = *A*//*T*_*δ*_(*A*), one can therefore conclude that *H*_*δ*_(*A*) is a non-trivial hierarchical classification of *A* and so, that ⟨*A, H*_*δ*_(*A*) ⟩ is well a finite non-trivial hierarchical classificatory structure. Hence our second (and more familiar) formalisation of *D*.

After that, consider the set

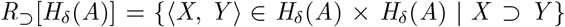

Given the extensional definition of *H*_*δ*_(*A*), one obtains

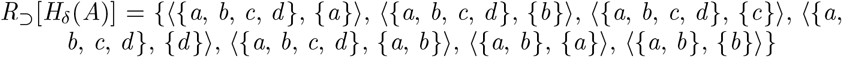

This set being a relation of hierarchical strict order on *H*_*δ*_(*A*), it follows that ⟨*H*_*δ*_(*A*), *R*_⊃_[*H*_*δ*_(*A*)] ⟩ is well a finite relational structure of hierarchical strict order. Hence our third formalisation of *D*. And finally, one can easily calculate the transitive reduction *R*_⊐_[*H*_*δ*_(*A*)] of *R*_⊃_[*H*_*δ*_(*A*)] by removing all the ordered pairs of the latter entailed by transitivity, i.e.,

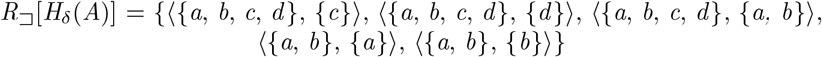

Such a set is a covering relation on *H*_*δ*_(*A*) and so, ⟨*H*_*δ*_(*A*), *R*_⊐_[*H*_*δ*_(*A*)] ⟩ is well a finite covering relational structure or, if you prefer, a hierarchical 4-tree such that its 4 leaves bijectively correspond to the 4 elements of *A*. Hence our fourth formalisation of *D*.

We now turn with the dendritic representations of ⟨*H*_*δ*_(*A*), *R*_⊃_[*H*_*δ*_(*A*)] ⟩ and ⟨*H*_*δ*_(*A*), *R*_⊐_[*H*_*δ*_(*A*)] ⟩. The first kind of dendritic representation that interest us in this paper is that of “arrow diagram”. Given a finite simple binary relational structure, i.e., an ordered pair ⟨*A, R*⟩ such that (i) *A* is finite non-empty set, and (ii) *R* is a binary relation on *A*, i.e., a set such that *R* ⊆ *A*^2^, its arrow diagram is a picture made up of dots and arrowed lines such that (iii) the dots bijectively represent the members of *A*, meaning that there is exactly one dot *f* (*x*) for any *x* ∈ *A*, and (iv) for all *x, y* ∈ *A*, there is exactly one arrowed line joining *f* (*x*) to *f* (*y*) if and only if ⟨*x, y*⟩ ∈ *R*. Figure 1a illustrates the arrow diagram of ⟨*H*_*δ*_(*A*), *R*_⊃_[*H*_*δ*_(*A*)] ⟩. Then, suppose that ⟨*A, R*⟩ is also a finite relational structure of strict order. In this case, its arrow diagram can be simplified into a “Hasse diagram” by the removing of all arrowed line corresponding to transitivity. In other words, the Hasse diagram of such an order is nothing but the arrow diagram of its corresponding covering relational structure. Figure 1b shows a diagram which is both the Hasse diagram of (*H*_*δ*_(*A*), *R*_⊃_[*H*_*δ*_(*A*)]) and the arrow diagram of ⟨*H*_*δ*_(*A*), *R*_⊐_[*H*_*δ*_(*A*)] ⟩. Finally, one can obtain a dendrogram from a given Hasse diagram for the specific case where the order ⟨*A, R*⟩ is also hierarchical from the following simplifications: (a) deletion of the dots representing the element of *A*, (b) replacement of the arrowed lines by non-arrowed lines, and (c) deletion of all the labels except those of the terminal dots. Note also that for the specific case where ⟨*A, R*⟩ represents the inclusive order of a given hierarchical classificatory structure ⟨*S, A*⟩, all the labels “{*x*}” (which represent the singletons of *A*) are replaced by labels “*x* “for all *x* ∈ *S*. Figure 1c depicts the dendrogram of both ⟨*H*_*δ*_(*A*), *R*_⊃_[*H*_*δ*_(*A*)] ⟩ and ⟨*H*_*δ*_(*A*), *R*_⊐_[*H*_*δ*_(*A*)] ⟩ that can be deduced from the arrow/Hasse diagram of figure 1b.

**Fig. 1.**
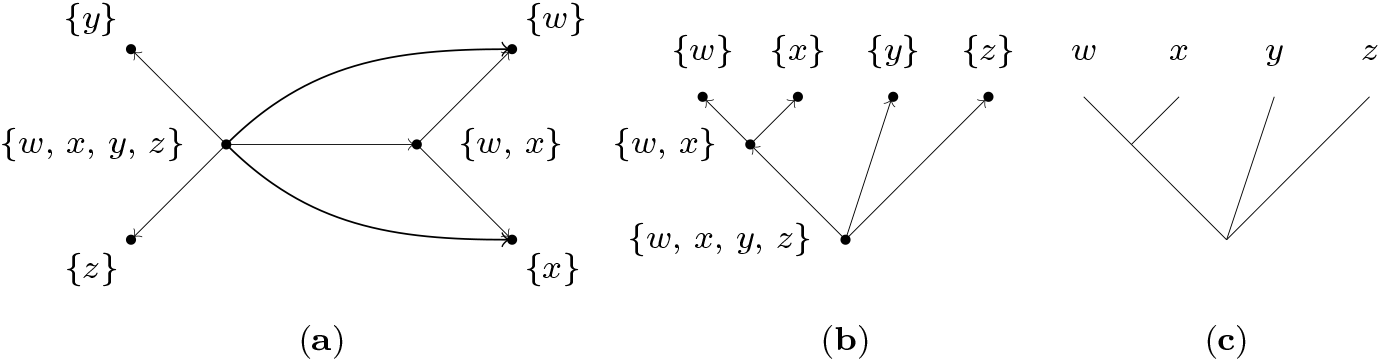
Dendritic representations of the order-like formalisations of the cladistic hypothesis *D* = ((*ab*)*cd*). **a** Arrow diagram of the relational structure of hierarchical strict order ⟨*H*_*δ*_(*A*), *R*_⊃_[*H*_*δ*_(*A*)] ⟩ such that *A* = {*a, b, c, d*}, *H*_*δ*_(*A*) = *H*_*A*0_ ∪ {{*a, b*}}, and *R*[*H*_*δ*_(*A*) = {⟨*X, Y* ⟩ ∈ *H*_*δ*_(*A*) × *H*_*δ*_(*A*) | *X* ⊃ *Y*}. **b** Arrow diagram of the corresponding covering relational structure (*H*_*δ*_(*A*), *R*_⊐_[*H*_*δ*_(*A*)]), i.e., Hasse diagram of ⟨*H*_*δ*_(*A*), *R*_⊃_[*H*_*δ*_(*A*)] ⟩. **c** Dendrogram of *D*; note that the latter can be obtained from the previous Hasse diagram from the following simplifications: deletion of the dots representing the clusters of *H*_*δ*_(*A*), replacement of the arrowed lines by non-arrowed lines, replacement of the label “{*x*}” by the label “*x* “for any *x* of *A*, and deletion of all the other labels.

## 6 Three-item statements

In phylogenetics, a 3is, also called a triad (Adams 1986: 302), a triplet (Vach 1994; Wilkinson 1994: 344), or still a rooted triple (Bryant 2003: 3) is an assertion saying that given three distinct objects (of the same kind), two of them are more closely related than either is to the third. More abstractly, that consists of saying that two of them are equivalent compared to the third (given a context or a parameter).

For example, a possible 3is for the three distinct elements *a, b*, and *c* of *A* is the one asserting that *a* and *b* are equivalent compared to *c*. If appropriate, one writes “(*ab*)*c*”, “*c*(*ab*)”, or still “*ab*|*c*”. Note that the order between *a* and *b* is irrelevant, i.e., (*ab*)*c* = (*ba*)*c*. Indeed, “*a* and *b* are equivalent compared to *c*” actually corresponds to the conjunction of “*a* is equivalent to *b* compared to *c*” and “*b* is equivalent to *a* compared to *c*”. But if true, then that also means for any *T* ∈ *𝒟*_*A*_ and all distinct *x, y, z* ∈ *A* that (*xy*)*z* is deductible from *T* if and only if {⟨*x, y, z* ⟩, ⟨*y, x, z* ⟩} ⊆ *T*.

Now let three distinct *x, y, z* ∈ *A*. In this case, the 3is (*xy*)*z* can be defined as follows:

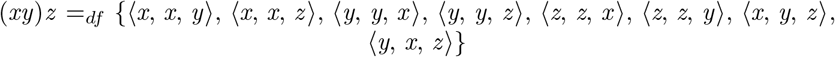

i.e.,

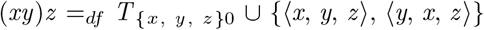

where *T* _{*x, y, z*}0_ denotes the trivial degree of equivalence relation on {*x, y, z*}. For any ⟨*T, H* ⟩ ∈ *𝒟*_*A*_ × *ℋ* _*A*_, and for all distinct *x, y, z* ∈ *A*, such a definition implies the following consequences: (i) (*xy*)*z* = (*yx*)*z*, (ii) for all distinct *u, v, w* ∈ *A*, (*uv*)*w* = (*xy*)*z* if and only if {*u, v*} = {*x, y*} and *w* = *z*, (iii) (*xy*)*z* is deductible from *T* if and only if (*xy*)*z* ⊆ *T* or, if you prefer, if and only if *z ∉* [*x* & *y*]_*T*_, (iv) (*xy*)*z* is deductible from *H* if and only if there is at least one *X* ∈ *I*_*H*_ such that {*x, y*} ⊆ *X* and *z ∉X*, and (v) the deduction of (*xy*)*z* from *T* is equivalent to the deduction of (*xy*)*z* from *H* if and only if *f* (*T*) = *H* and g(*H*) = *T*.

Then, suppose that *T* implies *k* 3is *t* _1_, *t* _2_, …, *t*_*k*_. In this case, one has (vi)

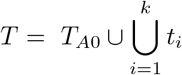

and (vii) either

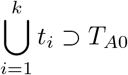

for the case where for all distinct *x, y* ∈ *A*, there are at least one 3is *t*_*i*_ ⊆ *T* and one *z* ∈ *A* such that *t*_*i*_ ∈ {(*xy*)*z*, (*xz*)*y*, (*yz*)*x*}, or

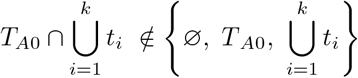

otherwise. In other words, the 3is deductible from *T* always correspond to the non-trivial part of the latter (*T* implies at least one 3is if and only if *T* ≠*T*_*A*0_), i.e.,

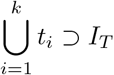

For the cladistic hypothesis *D* seen above, one can therefore write

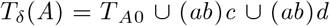

Now let ⟨*T, H* ⟩ ∈ *𝒟*_*A*_ × *ℋ* _*A*_, and let three distinct *x, y, z* ∈ *A*. In this case, (*xy*)*z* is part-whole compatible with *T* and/or *H* if and only if (*xy*)*z* is respectively deducible from *T* and/or *H*. Naturally, this implies that the part-whole compatibility of (*xy*)*z* with *T* is equivalent to the part-whole compatibility of (*xy*)*z* with *H* if and only if *f* (*T*) = *H* and *g*(*H*) = *T*. Then let (*uv*)*w* and (*xy*)*z* be two 3is. In this case, (*uv*)*w* and (*xy*)*z* are pairwise compatible if and only if there is at least one non-trivial degree of equivalence relation *T* on {*u, v, w*} ∪ {*x, y, z*} such that (*uv*)*w* and (*xy*)*z* are both part-whole compatible with *T*. But if true, then that also means that (*uv*)*w* and (*xy*)*z* are pairwise compatible if and only if there is at least one non-trivial hierarchical classification *H* of {*u, v, w*} ∪ {*x, y, z*} such that (*uv*)*w* and (*xy*)*z* are both part-whole compatible with *H*. It follows that (*uv*)*w* and (*xy*)*z* are pairwise incompatible if and only if {*u, v, w*} = {*x, y, z*} and (*uv*)*w* ≠ (*xy*)*z*. By contraposition, that also means that (*uv*)*w* and (*xy*)*z* are pairwise compatible if and only if either {*u, v, w*} {*x, y, z*} or (*uv*)*w* = (*xy*)*z*. Finally, let (*xy*)*z* be a 3is and let *T* ∈ *𝒟*_*A*_. In this case, (*xy*)*z* is compatible by aggregation with *T* if and only if there is at least one non-trivial degree of equivalence relation *T*_*a*_ on *A* ∪ {*x, y, z*} such that (i) *T*_*a*_ ⊇ *T* and (ii) (*xy*)*z* is part-whole compatible with *T*_*a*_. Thus, compatibility by aggregation generalises both part-whole compatibility and pairwise compatibility.

Another consequence of the previous definitions of 3is is that the properties seen above for any *T* ∈ *𝒟*_*A*_ can be rewritten in terms of 3is. For all distinct *w, x, y, z* ∈ *A*, this gives:

15. (*xy*)*z* ⊆ *T* → ¬[(*xz*)*y* ⊆ *T* ∨ (*yz*)*x* ⊆ *T*]

16. (*wx*)*z* ⊆ *T* → [(*wx*)*y* ⊆ *T* ∨ (*wy*)*z* ⊆ *T*]

17. [(*xy*)*w* ⊆ *T* ∧ (*yz*)*w* ⊆ *T*] → (*xz*)*w* ⊆ *T*

18. [(*wx*)*y* ⊆ *T* ∧ (*wy*)*z* ⊆ *T*] → [(*wx*)*z* ⊆ *T* ∧ (*xy*)*z* ⊆ *T*]

19. [(*wx*)*y* ⊆ *T* ∧ (*xy*)*z* ⊆ *T*] → [(*wx*)*z* ⊆ *T* ∧ (*wy*)*z* ⊆ *T*]

20. [(*wx*)*y* ⊆ *T* ∧ (*yz*)*w* ⊆ *T*] → [(*wx*)*z* ⊆ *T* ∧ (*yz*)*x* ⊆ *T*]

Among them, property 15, which is in fact another statement of the pairwise incompatibility property seen above, follows from properties 1 and 5. In the same way, property 16 is entailed by property 4, property 17 by property 3, property 18 by properties 9 or 10, property 19 by property 11, and property 20 translates property 12.

We can also write these properties less formally. Respectively, this gives (here too, for all distinct *w, x, y, z* ∈ *A*):

15. If (*xy*)*z*, then not (*xz*)*y* and not (*yz*)*x*

16. If (*wx*)*z*, then (*wx*)*y* and/or (*wy*)*z*

17. If (*xy*)*w* and (*yz*)*w*, then (*xz*)*w*

18. If (*wx*)*y* and (*wy*)*z*, then (*wx*)*z* and (*xy*)*z*

19. If (*wx*)*y* and (*xy*)*z*, then (*wx*)*z* and (*wy*)*z*

20. If (*wx*)*y* and (*yz*)*w*, then (*wx*)*z* and (*yz*)*x*

Note that properties 17 to 20 correspond to Dekker’s (1986) dyadic rules established for fully ramified undirected *n*-trees when they are reformulated for hierarchical *n*-trees by Wilkinson et al. (2004: 994) and completed by Rineau (2017) and Rineau et al. (2020). To be more precise, property 17 corresponds to the rooted variant of first Dekker’s dyadic rule, and properties 18 to 20 correspond to the rooted variants of the second one.

Now, consider the following writing rule: for all distinct *x, y, z* ∈ *A*, there is one ordered pair of parentheses “(“and “)” that contain *x* and *y* but not *z* – i.e., one may write “(… *x* … *y* …)… *z* … “where “… “stands for any other object of *A* – if and only if {⟨*x, y, z* ⟩, ⟨*y, x, z* ⟩} ⊆ *T*. In other words, any instance of “(… *x* … *y*…)… *z* … “means that there is at least one *X* ∈ *I*_*H*_ such that {*x, y*} ⊆ *X* and *z ∉ X*. In this case, one can rewrite properties 16 to 20 in terms of four-item statements (4is; Fig. 2), i.e., (still for any four distinct objects *w, x, y, z* ∈ *A*):

**Fig. 2.**
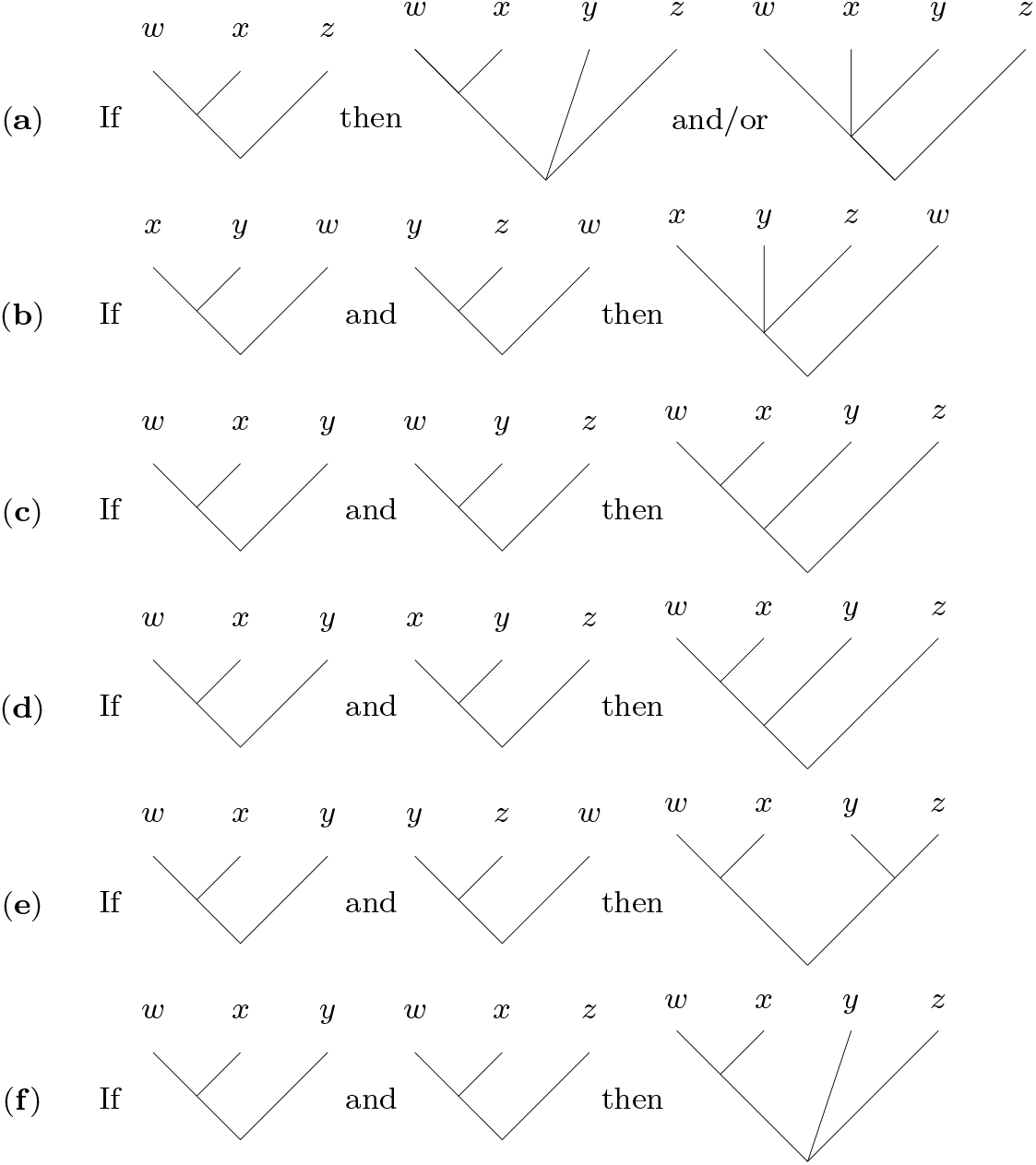
Illustration of 3is properties in terms of 4is: **a** insertion property, **b** in-out combination, **c** 1st asymmetric combination, **d** 2nd asymmetric combination, **e** symmetric combination, **f** in-in combination.

16. If (*wx*)*z*, then (*wx*)*yz* and/or (*wxy*)*z* (insertion property)

17. If (*xy*)*w* and (*yz*)*w*, then (*xyz*)*w* (in-out combination)

18. If (*wx*)*y* and (*wy*)*z*, then ((*wx*)*y*)*z* (1^*st*^ asymmetric combination)

19. If (*wx*)*y* and (*xy*)*z*, then ((*wx*)*y*)*z* (2^*nd*^ asymmetric combination)

20. If (*wx*)*y* and (*yz*)*w*, then (*wx*)(*yz*) (symmetric combination)

Let us make some words about the name of these properties that are given in parenthesis. On the one hand, property 16 is called “insertion property” because it actually translates the fact that there are seven possibilities for inserting the object *y* as a fourth leaf on the hierarchical 3-tree ((*wx*)*z*), i.e., (i) (((*wy*)*x*)*z*), (ii) ((*wxy*)*z*), (iii) ((*w* (*xy*))*z*), (iv) (((*wx*)*y*)*z*), (v) ((*wx*)*yz*), (vi) ((*wx*)(*yz*)), and (vii) (((*wx*)*z*)*y*). On the other hand, properties 17 to 20 are combination properties because they allow us to deduce a 4is from two 3is sharing two items. More precisely, property 17 is called “in-out combination” because of the distribution of these two shared items, and the names of the properties 18 to 20 follow from the shape of the obtained 4is.

Until here, all the combinatory relationships between 3is have been deduced only from basic and derived properties of degree of equivalence relations. It is however possible to deduce some additional properties by taking into consideration the mathematical equivalence between these relations and hierarchical classifications. For example, let ⟨*T, H* ⟩ ∈ *𝒟*_*A*_ × *ℋ* _*A*_ such that *H* = *f* (*T*) and *T* = *g*(*H*), and let four arbitrary distinct elements *w, x, y, z* ∈ *A*. Then, suppose that (*wx*)*y* ⊆ *T* and (*wx*)*z* ⊆ *T*. In this case, there are (at least) two clusters *X* and *Y* in *I*_*H*_ such that: (i) {*w, x*} ⊆ *X* and *y∉ X*, and (ii) {*w, x*} ⊆ *Y* and *z ∉Y*. Thus, *X* ∩ *Y* ⊇ {*w, x*} and so *X* ∩ *Y* ≠= ∅. But if true, then *X* ⊆ *Y* and/or *Y* ⊆ *X*, since *H* ∈ *ℋ* _*A*_. As a result, there is necessarily at least one *Z* ∈ *I*_*H*_ such that: (i) {*w, x*} ⊆ *Z* and {*y, z*} ∩ *Z* = ∅, and (ii) *Z* = *X* and/or *Z* = *Y*. By reconsidering the previous parenthesis-based symbolism, we can now write this property in terms of 4is:

21. If (*wx*)*y* and (*wx*)*z*, then (*wx*)*yz* (in-in combination)

As for property 17, its name follows from the distribution of the shared items between (*wx*)*y* and (*wx*)*z*.

## 7 Higher order rules

From these basic combination rules can be derived higher-order combination rules. In a non-exhaustive way, properties 17, 21, 18, and 19 can respectively be generalised as follows (Vautrin 2019):

22. If (*x*_1_*x*_2_)*y*, (*x*_2_*x*_3_)*y*, …, and (*x*_*k*−1_*x*_*k*_)*y*, then (*x*_1_*x*_2_…*x*_*k*_)*y* (generalised in-out combination)

23. If (*xy*)*z*_1_, (*xy*)*z*_2_, …, and (*xy*)*z*_*k*_, then (*xy*)*z*_1_*z*_2_…*z*_*k*_ (generalised in-in combination)

24. If (*x*_1_*x*_2_)*x*_3_, (*x*_1_*x*_3_)*x*_4_, …, and (*x*_1_*x*_*k*−1_)*x*_*k*_, then ((… (((*x*_1_*x*_2_)*x*_3_)*x*_4_)…)*x*_*k*−1_)*x*_*k*_ (generalised 1^*st*^ asymmetric combination)

25. If (*x*_1_*x*_2_)*x*_3_, (*x*_2_*x*_3_)*x*_4_, …, and (*x*_*k*−2_*x*_*k*−1_)*x*_*k*_, then (((… (((*x*_1_*x*_2_)*x*_3_)*x*_4_)…)*x*_*k*−2_)*x*_*k*−1_)*x*_*k*_ (generalised 2^*st*^ asymmetric combination)

Property 20 can be generalised in two main ways:

26. If (*x*_1_*x*_2_… *x*_*k*_)*y*_1_, (*x*_1_*x*_2_… *x*_*k*_)*y*_2_, …, (*x*_1_*x*_2_… *x*_*k*_)*y*_*n*_, and (*y*_1_*y*_2_… *y*_*n*_)*x*_1_, (*y*_1_*y*_2_… *y*_*n*_)*x*_2_, …,(*y*_1_*y*_2_… *y*_*n*_)*x*_*k*_, then (*x*_1_*x*_2_… *x*_*k*_)(*y*_1_*y*_2_… *y*_*n*_) (apical generalised symmetric combination)

27. If (*x*_1_*y*_1_)*x*_2_, (*x*_1_*y*_1_)*x*_3_, …, (*x*_1_*y*_1_)*x*_*k*_, and (*x*_2_*y*_2_)*x*_1_, (*x*_2_*y*_2_)*x*_3_, …, (*x*_2_*y*_2_)*x*_*k*_ and (*x*_*k*_*y*_*k*_)*x*_1_, (*x*_*k*_*y*_*k*_)*x*_2_, …, (*x*_*k*_*y*_*k*_)*x*_*k*−1_, then (*x*_1_*y*_1_)(*x*_2_*y*_2_)… (*x*_*k*_*y*_*k*_) (basal generalised symmetric combination)

To give an idea of the range of rules that can be inferred from the 3is rules, one can also mix the previous rules. Let us consider for example two compatible trees with only one informative node having a common cluster (including the same terminals). If the common cluster is the root, then there are three possibilities:

22. If (*x*_1_*x*_2_… *x*_*k*_)*x*_*k*+1_… *x*_*l*_ and (*x*_*k*+1_… *x*_*l*_)*x*_1_*x*_2_… *x*_*k*_, then (*x*_1_*x*_2_… *x*_*k*_)(*x*_*k*+1_… *x*_*l*_)

23. If (*x*_1_*x*_2_… *x*_*k*_)*x*_*k*+1_… *x*_*m*_ and (*x*_*k*+1_… *x*_*l*_)*x*_1_*x*_2_… *x*_*k*_*x*_*l*+1_… *x*_*m*_, then (*x*_1_*x*_2_… *x*_*k*_)(*x*_*k*+1_… *x*_*l*_)*x*_*l*+1_… *x*_*m*_

24. If (*x*_1_*x*_2_… *x*_*k*_*x*_*k*+1_… *x*_*l*_)*x*_*l*+1_… *x*_*m*_ and (*x*_1_*x*_2_… *x*_*k*_)*x*_*k*+1_… *x*_*m*_, then ((*x*_1_*x*_2_… *x*_*k*_)*x*_*k*+1_… *x*_*l*_)*x*_*l*+1_… *x*_*m*_

If the common cluster is the root for one tree but not for the other, then there is only one possibility:

22. If (*x*_1_*x*_2_… *x*_*k*_*x*_*k*+1_… *x*_*l*_)*x*_*l*+1_… *x*_*m*_ and (*x*_1_*x*_2_… *x*_*k*_)*x*_*k*+1_… *x*_*l*_, then ((*x*_1_*x*_2_… *x*_*k*_)*x*_*k*+1_… *x*_*l*_)*x*_*l*+1_… *x*_*m*_

If the common cluster is not the root in both trees, then there is only one possibility:

22. If (*x*_1_*x*_2_… *x*_*k*_)*x*_*k*+1_… *x*_*l*_ and (*x*_1_*x*_2_… *x*_*k*_)*x*_*l*+1_… *x*_*m*_, then (*x*_1_*x*_2_… *x*_*k*_)*x*_*k*+1_… *x*_*l*_*x*_*l*+1_… *x*_*m*_

The set of these higher-order rules allows the following definition, namely that any pair of trees sharing a cluster in common (by all the terminals it contains) can be unambiguously combined into a tree.

## 8 Application

Bryant and Steel (1995: 448, theorem 3) have shown that there is an infinite number of 3is *k*-adic rules irreducible to *l*-adic rules such that 2 *l < k* for any integer *k >* 3. Based on an example of five compatible but pairwise non-combinable 3is, we show here how it is possible, at least for this example, to reduce this *5*-adic rule to the 3is properties stated previously.

To simplify the reading, we will not detail all of the 3is generated by the combinations, but only the intermediate trees generated by the inference rules (thus, a combination can take up 3is from a tree previously inferred). Moreover, higher-order inference rules stated in the above part are also used for simplicity. Let {*a, b, c, d, e, f, g*} be a set of seven elements and {(*ab*)*g*, (*dg*)*e*, (*cd*)*a*, (*df*)*b*, (*ef*)*a*} be a set of five triplets (Fig. 3a; Wilkinson pers. com.). Let us start from the following property:

**Fig. 3.**
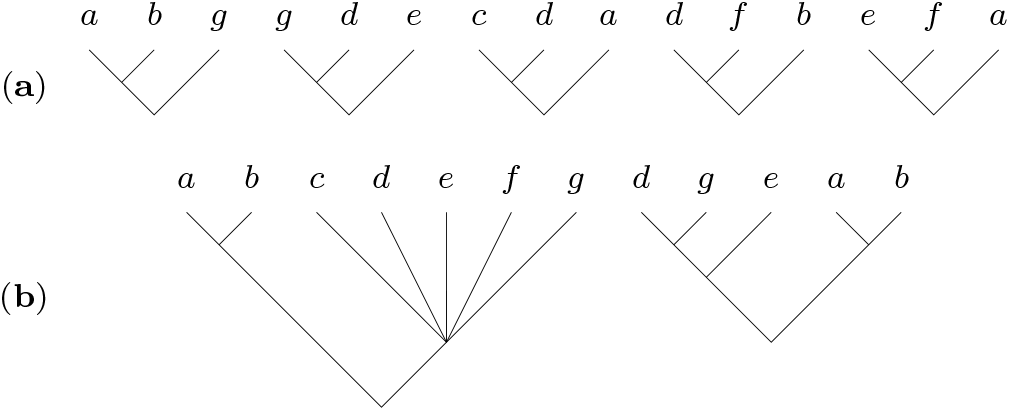
Illustration of the set of five 3is non-pairwise combinable by application of basic dyadic combination properties, but combinable as a whole into larger trees. **a** the five initial 3is, **b** the two cladistic trees implied by them.

– If (*dg*)*e* then (*adg*)*e* and/or (*dg*)*ae* (2-3 negative transitivity)

We begin by demonstrating that knowing {(*ab*)*g*, (*dg*)*e*, (*cd*)*a*, (*df*)*b*, (*ef*)*a*}, then (*adg*)*e* is incompatible with them. If (*adg*)*e* is not possible, then (*dg*)*ae* is necessarily true.

### 8.1 If (*gda*)*e* is true, then

– If (*ad*)*e* and (*cd*)*a*, then ((*cd*)*a*)*e* (asymmetric combination)
– If (*ad*)*e* and (*ef*)*a*, then (*ad*)(*ef*) (symmetric combination)
– If (*cd*)*a* and (*ad*)*f*, then ((*cd*)*a*)*f* (asymmetric combination)
– If (*ef*)*a* and (*ac*)*f*, then (*ef*)(*ac*) (symmetric combination)
– If (*cd*)*a* and (*cd*)*e* and (*cd*)*f*, then (*cd*)*aef* (generalized in-in combination)
– If (*ef*)*a* and (*ef*)*c* and (*ef*)*d*, then (*ef*)*acd* (generalized in-in combination)
– If (*cda*)*e* and (*cda*)*f*, then (*cda*)*ef* (combination by common node)
– If (*cd*)*aef* and (*cda*)*ef* and (*ef*)*acd*, then ((*cd*)*a*)(*ef*) (combination by common node)

*addition of g*

– If (*ef*)*d* and (*dg*)*e*, then (*ef*)(*gd*) (symmetric combination)
– If (*cd*)e and (*dg*)*e*, then (*cdg*)*e* (in-out combination)
– If (*ef*)*g* and (*cg*)*e*, then (*ef*)(*cg*) (symmetric combination)
– If (*ef*)a*g* and (*ef*)*c* and (*ef*)*d* and (*ef*)*g*, then (*ef*)*acdg* (generalized in-in combination)
– If (*cg*)*f* and (*ad*)*f* and (*cd*)*f*, then (*acdg*)*f* (generalized in-out combination)
– If (*dg*)*e* and (*ad*)*e* and (*cd*)*e*, then (*acdg*)*e* (generalized in-out combination)
– If (*acdg*)*e* and (*acdg*)*f*, then (*acdg*)*ef* (combination by common node)
– If (*acdg*)*ef* and (*ef*)*acdg*, then (*ef*)(*acdg*) (combination by common node)

*addition of b*

– If (*ab*)*g* and (*ag*)*e*, then ((*ab*)*g*)*e* (asymmetric combination)
– If (*ab*)*g* and (*ag*)*f*, then ((*ab*)*g*)*f* (asymmetric combination)
– If (*ab*)*e* and (*ac*)*e*, then (*abc*)*e* (in-out combination)
– If (*ad*)*f* and (*ab*)*f*, then (*abd*)*f* (in-out combination)

But (*abd*)*f* implies (*bd*)*f*, which contradicts (*df*)*b* (incompatibility property). Thus (*adg*)*e* is false and so, (*dg*)*ae* is necessarily true (disjunctive syllogism).

### 8.2 If (*dg*)*a* is true, then

– If (*dg*)*a* and (*ab*)*g*, then (*dg*)(*ab*) (symmetric combination)
– If (*dg*)*a* and (*cd*)*a*, then (*cdg*)*a* (in-out combination)
– If (*cd*)*a* and (*ab*)*d*, then (*ab*)(*cd*) (symmetric combination)
– If (*dg*)*b* and (*cd*)*b*, then (*cdg*)*b* (in-out combination)
– If (*cdg*)*a* and (*gcd*)*b*, then (*cdg*)*ab* (combination by common node)
– If (*ab*)*c* and (*ab*)*d* and (*ab*)*g*, then (*ab*)*cdg* (generalized in-in combination)
– If (*cdg*)*a* and (*ab*)*cdg*, then (*cdg*)(*ab*) (combination by common node)

*addition of f*

– If (*df*)*b* and (*ab*)*d*, then (*ab*)(*df*) (symmetric combination)
– If (*df*)*a* and (*dg*)*a*, then (*dfg*)*a* (in-out combination)
– If (*df*)*b* and (*dg*)*b*, then (*dfg*)*b* (in-out combination)
– If (*df*)*b* and (*dg*)*b* and (*cd*)*b*, then (*cdfg*)*b* (combination by common node)
– If (*df*)*a* and (*dg*)*a* and (*cd*)*a*, then (*cdfg*)*a* (combination by common node)
– If (*cdfg*)*a* and (*cdfg*)*b*, then (*cdfg*)*ab* (combination by common node)
– If (*ab*)*c* and (*ab*)*d* and (*ab*)*f* and (*ab*)*g, then* (*ab*)*cdfg* (generalized in-in combination)
– If (*cdfg*)*ab* and (*ab*)*cdfg*, then (*ab*)(*cdfg*) (combination by common node)

*addition of e*

– If (*ef*)*a* and (*ab*)*f*, then (*ab*)(*ef*) (symmetric combination)
– If (*df*)*b* and (*dg*)*b* and (*cd*)*b* and (*ef*)*b*, then (*cdefg*)*b* (generalized in-in combination)
– If (*df*)*a* and (*dg*)*a* and (*cd*)*a* and (*ef*)*a*, then (*cdefg*)*a* (generalized in-in combination)
– If (*cdefg*)*a* and (*cdefg*)*b*, then (*cdefg*)*ab* (combination by common node)
– If (*ab*)*c* and (*ab*)*d* and (*ab*)*e* and (*ab*)*f* and (*ab*)*g*, then (*ab*)*cdefg* (generalized in-in combination)
– If (*dfgcfe*)*ab* and (*ab*)*cdefg*, then (*ab*)(*cdefg*) (combination by common node)
– If (*gd*)*e* and (*eg*)*a*, then ((*gd*)*e*)*a* (asymmetric combination)
– If (*gd*)*e* and (*eg*)*b*, then ((*gd*)*e*)*b* (asymmetric combination)
– If ((*gd*)*e*)*a* and ((*gd*)*e*)*b*, then ((*gd*)*e*)*ab* (combination by common node)
– If ((*gd*)*e*)*ab* and (*ab*)*gde*, then ((*gd*)*e*)(*ab*) (combination by common node)

In summary, if (*ab*)*g*, (*gd*)*e*, (*cd*)*a*, (*df*)*b*, and (*ef*)*a*, then (*ab*)(*cdefg*) and ((*dg*)*e*)(*ab*) (Fig. 3).

Consequently, only a finite and few numbers of primary inference rules are needed, all of the other being deductible. One can see that the key property allowing this is the 2-3 negative transitivity. The same result can be obtained empirically by coding the five 3is in a parsimony matrix (Nelson and Platnick 1991) and resuming the obtained optimal trees using a reduced cladistic consensus (Wilkinson 1994). Such a result suggests that using the 2-3 negative transitivity property could allow us to reduce all *k*-adic rules of combination for 3is to a finite set of more basic rules. Given the strong connexion between rooted trees and unrooted trees (Bryant and Steel 1995: 448-449), one can expect that our proposal for 3is is transposable to quartets.

## Acknowledgements

We thank Mark Wilkinson for sharing with us a stimulating example of five three-item statements, and Mike Steel for sending us Dekker’s remarkable dissertation. We are also grateful to Paul Zaharias for his helpful comments on the manuscript. Finally, we thank Rachel Vautrin, Renée Zaragüeta, Paul Zaharias, and all the members of the Paris working group on 3ia and Cladistics for fruitfull discussions.

## Footnotes

1 Given a phylogenetic/cluster analysis, a TU is any taxon defined prior to this analysis and taken as an object by this analysis. Usually, phylogenetic analyses seek to establish kinship relationships of this or that kind (e.g., the ancestor-descendant relationship or the degree of kinship relationship) between several mutually disjoint TU. These taxa can be contrasted to hypothetical taxa (HT), i.e., taxa highlighted by phylogenetic reconstruction (e.g., inclusive clades in cladistic phylogenetics).

2 For previous axiomatic approaches on these relations, see Colonius and Schulze (1981) and McMorris and Powers (2003).

3 Although some authors also use this term in biology (e.g., Ridley 2004: 472), these classifications are primarily known in mathematics as total hierarchies (Benzecri 1972: 30), hierarchies (Leclerc 1985a, 1985b; Vach 1994: 61; McMorris and Powers 2003: 42), hierarchies of parts (Barthélemy and Guénoche 1991: 22-23), or even *n*-trees for the specific case where *A* is finite (Margush and McMorris 1981: 240; Adams 1986: 304; McMorris and Powers 2003: 42). A related (and more general) concept is that of tree of subsets sensu Estabrook and McMorris (1980: 369). See Leclerc (1985a, 1985b) for an extensive study of them.

4 This formalisation should be understood as a minimal one. Indeed, phylogenetic hypotheses can be more complex by the incorporation of other kinds of information (e.g., branch lengths for tree-like and network-like phylogenetic hypotheses). However, even such complex phylogenetic hypotheses always involve a finite non-empty set of (biological) objects and a phylogenetic relationship (of a given arity) on them.

